# Dissecting transcriptomic signatures of genotype x genotype interactions during the initiation of plant-rhizobium symbiosis

**DOI:** 10.1101/2020.06.15.152710

**Authors:** Camilla Fagorzi, Giovanni Bacci, Rui Huang, Lisa Cangioli, Alice Checcucci, Margherita Fini, Elena Perrin, Chiara Natali, George Colin diCenzo, Alessio Mengoni

**Author notes:** Department of Agricultural and Food Science, University of Bologna, Bologna, Italy (Alice Checcucci).

## Abstract

Rhizobia are ecologically important, facultative plant symbiotic microbes. In nature there exists large variability in the association of rhizobial strains and host plant of the same species. Here, we evaluated whether plant and rhizobial genotypes influence the initial transcriptional response of rhizobium following perception of host plant. RNA-sequencing of the model rhizobium *Sinorhizobium meliloti* exposed to root exudates or luteolin was performed in a combination of three *S. meliloti* strains and three *Medicago sativa* varieties. The response to root exudates involved hundreds of changes in the rhizobium transcriptome. Of the differentially expressed genes, expression of 35% were influenced by strain genotype, 16% by the plant genotype, and 29% by strain x host plant genotype interactions. We also examined the response of a hybrid *S. meliloti* strain, in which the symbiotic megaplasmid (~ 20% of the genome) was mobilized between two of the above-mentioned strains. Dozens of genes resulted up-regulated in the hybrid strain, indicative of nonadditive variation in the transcriptome. In conclusion, this study demonstrated that transcriptional responses of rhizobia upon perception of legumes is influenced by the genotypes of both symbiotic partners, and their interaction, suggesting a wide genetic spectrum of partner choice selection in plant-rhizobium symbiosis.

## Introduction

Microbes play a crucial role in the biology and evolution of their eukaryotic hosts [1]. Among other activities, microbes contribute to the host’s acquisition of nutrients [2], functioning of the host’s immune system [3], and protection of the host from predation [4]. The rules governing host-microbe interactions remain a topic of intense investigation. In many cases, the eukaryotic host selectively recruits the desired microbial partner; squid light organs are selectively colonized by *Vibrio* symbionts [5], legumes select for effective symbionts by sanctioning non-effective symbionts [6], and the crop microbiome is cultivar-dependent [7, 8]. The genetic basis determining the quality of a microbial symbiont and its ability to effectively colonize its eukaryotic partner is generally not well-understood, but high-throughput genome sequencing projects of host-associated microbes and complete microbiomes are shedding light on this topic [9–12]. In the case of plants, such studies have observed an enrichment of certain gene functions in plant-associated microbes, such as genes related to carbohydrate metabolism, secretion systems, phytohormone production, and phosphorous solubilization [11–14].

The rhizobia are an ecologically important exemplar of facultative host-associated microbes. These soil-dwelling bacteria are able to colonize plants and enter into an endosymbiotic association with plants of the family *Fabaceae* [15]. This developmentally complex process begins with an exchange of signals between the free-living organisms [16], which leads to invasion of the plant by the rhizobia [17], and culminates in the formation of a new organ (a nodule) in which the plant cells are intra-cellularly colonized by N_2_-fixing rhizobia [18, 19]. Decades of research have identified an intricate network of coordinated gene functions required to establish a successful mutualistic interaction between rhizobia and legumes [19–21]. In contrast to the core symbiotic machinery, most of which has been elucidated, much remains unknown about the accessory genes required to optimize the interaction.

In addition to simple gene presence/absence, genotype by genotype (GxG) interactions have prominent impacts on symbiotic outcome [22]. The importance of both the plant and bacterial genotypes, and their interaction, in optimizing symbioses between rhizobia and legumes was recognized in early population genetic studies [23–25]. More recently, greenhouse studies have directly demonstrated the influence GxG interactions on the fitness of both the plant and rhizobium partners [26–29]. The newly developed select-and-resequence approach is providing a high-throughput approach to begin uncovering the genetic basis underlying GxG interactions on fitness in rhizobium – legume symbioses, as well as a way to screen for strain-specific effects of individual genes [30, 31]. To date, GxG interaction studies have largely focused on measurements of fitness as a holistic measure of the entire symbiotic process. Nodule formation is a complex developmental process involving several steps that each require a distinct molecular toolkit [32], and in principle, distinct GxG interactions could be acting at each of these developmental stages. Transcriptomic studies have demonstrated that GxG interactions have significant impacts on the gene expression patterns of both partners in mature N_2_-fixing nodules [33, 34]. However, we are unaware of studies specifically focusing on the role of GxG interactions in early developmental stages, such as during the initial perception of the partners by each other. Such knowledge is critical not only to fully understand the microevolution of host-associated bacteria, but also to develop improved rhizobium bioinoculants able to outcompete the indigenous rhizobium population [35, 36].

Here, we evaluated whether GxG interactions could be identified in the initial transcriptional response of rhizobium perception of a host plant. We worked with *Sinorhizobium meliloti*, which is one of the best studied models for GxG interactions in rhizobia. *S. meliloti* forms N_2_-fixing nodules on plants belonging to the tribe *Trigonelleae* [37] that includes *Medicago sativa* (alfalfa), a major forage crop grown worldwide for which many varieties have been developed [38]. The *S. meliloti* genome comprises three main replicons, a chromosome, a chromid, and a megaplasmid; the latter one harbours most of the essential symbiotic functions, including the genes responsible for the initial molecular dialogue with the host plant (*nod* genes) [39, 40]. To address our aim, the gene expression patterns of three strains of *S. meliloti* (each with distinct symbiotic properties) following four hours of exposure to root exudates derived from three *M. sativa* varieties were characterized using RNA-sequencing. Additionally, the relevance of the megaplasmid in defining the strain-specific transcriptional responses was analysed through studying a hybrid *S. meliloti* strain, in which the native megaplasmid was replaced with that of another wild type strain. The results demonstrated that the transcriptional response involved genes on all three replicons and that, even among conserved *S. meliloti* genes, transcriptional patterns were both strain and root exudate specific.

## Materials and Methods

### Strains and microbiological methods

The list of strains, and their host plant of origin, is reported in **Table S1**. *S. meliloti* Rm1021 is a spontaneous streptomycin-resistant derivative of the isolate SU47 recovered form *M. sativa* root nodules [41]. *S. meliloti* BL225C was isolated by plant trapping with the *M. sativa* variety “Lodi” in Lodi, Italy in 1996 [24]. *S. meliloti* AK83 strain was isolated from nodules of *M. falcata* grown in soil samples from the North Aral Sea region in Kazakhstan in 2001 [42]. *S. meliloti* BM806 (later termed as “hybrid strain”) is a Rm2011 (a near identical strain to Rm1021, as both are independent streptomycin resistant derivatives of the nodule isolate *S. meliloti* SU47 [43]) derivative in which the pSymA megaplasmid was replaced with the homologous megaplasmid (pSINMEB01) from strain BL225C [44]. Strains were grown at 30°C in TY with 0.2 g/l CaCl_2_, or in M9 supplemented with 0.2% succinate as the carbon source. For Rm1021, streptomycin (200 μg/mL) was added to the culture medium during routine growth.

### Plant varieties, root adhesion tests, and symbiotic assays

Three plant varieties were used (**Table S1**), differing in fall dormancy and in genotype. Fall dormancy (FD) is an important trait having large impacts on the productivity and persistence of alfalfa [45]. Cultivars Camporegio and Verbena are included in the subgroup of fall dormant type (FDT; FD 1–4), while cultivar Lodi is a semi-dormant type (SDT; FD 5–7). Symbiotic assays were performed, as previously reported [46], on 12 plants per strain – cultivar combination. The root adhesion test was performed five days following the inoculum of plantlets. Using sterile tweezers, plantlets were carefully removed from the substrate and divided into epicotyl and hypocotyl (i.e. root) portions. After measuring their length, roots were washed to remove loosely adherent cells by vortexing for ten seconds in 500 μl of 0.9% NaCl. Then, roots were transferred to 500 μl of fresh 0.9% NaCl, and vortexed for 30 seconds to collect bacterial cells strongly adhered to the root surface. Roots were removed from the tube and the quantity of bacterial cells detached from the roots and recovered in the NaCl solution was evaluated using Real Time PCR (qPCR) by a standard curve method on the *nodB* gene in a QuantStudio™ 7 flex (Applied Biosystems), as previously described [47, 48]. Differences were evaluated by one-way ANOVA Tukey pairwise contrast and using the Scott-Knott procedure as implemented in R [49].

### Root exudate production and metabolomic analyses

Root exudates were produced as previously reported [50] in two independent experiments, giving two biological replicas for each plant cultivar. A blank sample was prepared with the same setup of the plant experiment, but without adding the plantlet. Elemental analysis (CHNS) was performed on crude root exudates (a combined sample for each cultivar) using a carbon hydrogen and nitrogen analyzer (CHN-S Flash E1112, Thermofinnigan, San José, California, United States). For LC-MS, the extraction of the seven samples (two biological replicates per cultivar, plus the blank) was performed by metaSysX GmbH (www.metasysx.com) with a modified protocol from [51]. The samples were measured with a Waters ACQUITY Reversed Phase Ultra Performance Liquid Chromatography (RP-UPLC) coupled to a Thermo-Fisher Q-Exactive mass spectrometer that consists of an ElectroSpray Ionization source (ESI) and an Orbitrap mass analyzer (UPLC-MS). A C18 column was used for the chromatographic separation of the hydrophilic compounds. The mass spectra were acquired in full scan MS positive and negative modes (Mass Range [100−1500]). Extraction of the LC-MS data was accomplished with the software REFINER MS® 10.5 (GeneData, genedata.com). After extraction of the peak list from the chromatograms, data were processed, aligned, and filtered using in-house software. Only those features (peak IDs) that were present in at least two out of the seven samples were kept. At this stage, an average retention time (RT) and average m/z values were given to each feature. The alignment was performed for each platform independently (polar phase positive mode, polar phase negative mode). The annotation of the content of the sample was accomplished by matching the extracted data from the chromatograms with metaSysX’s library of reference compounds. Data from both platforms (RP-UPLC and UPLC-MS) were combined to build the final data matrix. Attribution of brute formulas to known compound was done using the PubChem database (pubchem.ncbi.nlm.nih.gov/). Principal Component Analysis (PCA) was performed on the Bray-Curtis dissimilarity obtained from each peak ID value. Statistical differences in single metabolites were assessed by Simper analysis based on the decomposition of the Bray-Curtis dissimilarity obtained from each peak ID value. All statistical analyses were done with the vegan package of R [52].

### RNA isolation

Overnight cultures of *S. meliloti*, grown in M9-succinate medium at 30°C at 130 rpm, were diluted to an OD_600_ of 0.05 in 5 ml of M9-succinate and incubated until an OD_600_ of 0.4 was reached. Then, either 10 μM of luteolin (Sigma-Aldrich) or one of the alfalfa root exudate (normalized by the total organic carbon as measured by the CHNS analysis; 0.250 ml, 0.042 ml, 0.224 ml, and 0.806 ml for Camporegio, Lodi, Verbena, and the blank samples, respectively) was added to each of the cultures and incubated for a additional 4 hours at 30°C with shaking at 130 rpm. Biological replicates were performed for each of the three strains across the five conditions. After incubation, cells were blocked with RNAprotect Bacteria (Qiagen, Venlo, The Netherlands) and total RNA was extracted using RNeasy Mini kits (Qiagen) from 0.5 ml of culture following the manufacturer’s instructions, including on column DNase I treatment. After elution, a second DNase I (ThermoFisher, Waltham, Massachusetts, USA) treatment was performed. The absence of contaminant DNA was verified by qPCR on the *nodC* gene of *S. meliloti*. Quality and quantity of extracted RNA were checked by spectrophometric readings (NanoQuant plate, Infinite PRO 200, Tecan, Männedorf, Switzerland), fluorometric measurement (Qubit, ThermoFisher), and cartridge electrophoresis on a 2100 Bioanalyzer (Agilent RNA Nano kit 6000, Agilent Technology, Santa Clara, California, USA). All RNA samples gave RNA Integrity Number (RIN) values between 9 and 10.

### Reverse transcriptase qPCR

Single stranded cDNA libraries were prepared from total RNA samples using SuperScript II reverse transcriptase (ThermoFisher) following the manufacturer’s instructions. qPCR was performed using a QuantStudio™ 7 flex (Applied Biosystems, Foster City, California, USA) programmed with the following temperature profile: 2 min at 94°C, followed by 40 cycles composed of 15 s at 94°C, 15 s at 60°C, and 30 s at 72°C, with a final melting curve to check for product specificity. Technical triplicate were carried-out as described in [48]. Gene *smc01804* (*rplM*), encoding the 50S ribosomal protein L13, was used as a housekeeping gene for normalization of the expression data. The list of primers used is reported in **Table S2**. Relative quantification (RQ, as 2-ΔΔLt) values were calculated with the ExpressionSuite ver. 1.0.4 software (Applied Biosystems). Differences on RQ data were evaluated by one-way ANOVA with Tukey pairwise contrast in R [49].

### RNA-sequencing and data analysis

Ribosomal RNA depletion was performed using MICROBexpress kits (ThermoFisher) following the manufacturer’s instruction starting from 0.6-1 μg of total RNA per sample. Removal of rRNA was checked on a Bioanalyzer 2100 (Agilent RNA Nano kit 6000, Agilent Technology). Ribosomal RNA depleted RNA preparations were used for library construction with TruSeq Stranded Total RNA Library Prep Gold kit (Illumina, San Diego, California, USA), using SuperScript II reverse transcriptase (ThermoFisher) for cDNA preparation. Libraries were assessed for quality using a DNA 1000 chip on a Bioanalyzer 2100 (Agilent Technologies), running 1 μl of each undiluted DNA library. Library normalization was performed based on Qubit fluorometric quantification. Libraries were sequenced on an Illumina Novaseq 6000 apparatus with a SP flow cell.

### Read mapping, counting, and differential expression analysis

Reads were demultiplexed using “bcl2fastq2” version 2.2 with default parameters. Demultiplexed sequences were then quality controlled using the StreamingTrim algorithm (version 1.0) [53] with a quality threshold of 20 Phred. Reads were mapped back to transcripts using Salmon (version 1.1.0) [54] against *decoy-aware* datasets containing both cDNA and the genome of each strain (as described in Salmon documentation: https://salmon.readthedocs.io/en/latest/salmon.html#preparing-transcriptome-indices-mapping-based-mode). Quantification files produced by Salmon were then imported into R using tximport package (version 1.10.1) [55]. Differential abundance analysis was performed with DESeq2 package (version 1.22.2) [56] on single strains in different conditions. To analyze all strains together, transcripts were collapsed into orthologous groups with Roary [57], version 3.11.2 (for additional details see the subchapter below). Counts produced by Salmon were collapsed following the group ID provided by Roary, producing a single table with ortholog-level quantification of transcripts. The produced table was then used to perform nested likelihood ratio test (LRT) with DESeq2. Strains and conditions were used together with their interaction to build a model for each group. Terms were then removed one by one to test their impact on the likelihood of the full model (as described in the DESeq2 documentation: http://bioconductor.org/packages/devel/bioc/vignettes/DESeq2/inst/doc/DESeq2.html#likelihood-ratio-test).

### Statistical analysis of differentially expressed genes

For each *S. meliloti* strain, genes differentially expressed (log_2_[fold change] ≥ 1, *p*-value < 0.01) in at least one condition (exposure to luteolin, or *M. sativa* Camporegio, Verbena, or Lodi root exudate) relative to the control condition were identified, and all fold change values (relative to the control) for these genes were extracted. Heatmaps of the differentially expressed genes (DEGs) were prepared for each strain using the *ComplexHeatmap* and *Heatmaply* packages of R [58, 59].

To compare expression of genes conserved between Rm1021, AK83, and BL225C, the pangenome of the three strains was calculated using Roary ver. 3.13.0 [57] with an identity threshold of 90%, and the genes found in all three strains (the core genes) were recorded. For each condition, core genes differentially expressed in at least one strain relative to the control condition were identified, and the fold change values for the gene and its orthologs in the other strains were extracted. Heatmaps of the differentially expressed genes (DEGs) were prepared for each strain using the *ComplexHeatmap* and *Heatmaply* packages of R. In addition, all fold change values for all of the core *S. meliloti* genes were extracted and used to run a PCA using the *prcomp* function of R that was visualized with the *ggplot2* package of R [60]. The same approach was used to compare expression of core genes across Rm1021, BL225C, and the hybrid strain.

All genes of *S. meliloti* strains Rm1021, AK83, and BL225C were functionally annotated using the standalone ver. 2 of eggNOG-mapper [61, 62] using default settings with the following two modifications: mode was set to diamond and query-cover was set to 20. Next, the Kyoto Encyclopedia of Genes and Genomes (KEGG) module annotations for each gene were extracted; if a gene was annotated with multiple KEGG modules, only the first one was kept. Then, for each strain-condition pairing, KEGG modules that were over- or under-represented among the up- and down-regulated genes (relative to the whole genome) were identified using hypergeometic tests (*p*-value < 0.05) with a custom R script. The same procedure was used to identify Cluster of Orthologous Genes (COG) categories that were over- or under-represented among the up- and down-regulated genes.

All data processing was performed using custom Python scripts using Python ver. 3.6.9 and the external libraries Pandas ver. 0.23.4 [63], pickle ver. 4.0, and numpy ver. 1.15.4 [64].

For each DEG, nested likelihood ratio tests (LRT) was used to evaluate the statistical significance of strain, condition (luteolin and treatments with root exudates from the three cultivar) and strain x condition interaction effects on gene expression. Implementation of nested LRT was performed as in [34], using a custom R script.

### Data availability

Gene expression data are available at GEO under the accession: Custom scripts developed for this work can be found in the GitHub repository: https://github.com/hyhy8181994/Sinorhizobium-RNAseq-2020.

## Results

### Symbiotic phenotypes differ across rhizobial strain x plant variety combinations

Symbiotic phenotypes (plant growth, nodule number) and root adhesion of *S. meliloti* strains Rm1021, BL225C, and AK83 were measured during interaction with three varieties of alfalfa (Camporegio, Verbena, Lodi). The results indicated that these phenotypes are influenced by both the plant and the bacterial genotypes (**Figure 1, Supplemental file S1**). Root adhesion phenotypes (**Figure 1a**) were divided by Scott-Knott test into three main groups reflecting high, medium, and low root colonization. Interestingly, each group was heterogenous with respect to both plant variety and *S. meliloti* strain, consistent with a specificity of plant variety (i.e. genotype “sensu lato”) and strain individuality (i.e. strain genotype) pairs in root colonization efficiency. For instance, *S. meliloti* BL225C strongly colonized the roots of the Camporegio and Verbena varieties, but it displayed much weaker colonization of the Lodi cultivar. On the other hand, *S. meliloti* AK83 colonized the Lodi and Camporegio varieties better than the Verbena cultivar. Nodules per plant, as well as measures of symbiotic efficiency (epicotyl length and the shoot dry weight), showed differences among the strain-variety combinations (**Figure 1b-d**). However, the extents of the variation were lower than those recorded for plant root adhesion. The highest number of nodules was found on the Lodi variety nodulated by *S. meliloti* AK83, which was previously interpreted as a consequence of its reduced N_2_-fixation ability with some alfalfa varieties [42, 48, 65]. Interestingly, the measures of symbiotic efficiency did not correlate with root adhesion phenotypes (both adhesion vs. dry weight and adhesion vs. epicotyl length gave not significant value of Pearson correlation, p >0.18). For example, the largest plants were the Lodi variety inoculated with *S. meliloti* BL225C despite the root adhesion of this combination being the lowest. Similarly, the smallest plants were the Verbena variety inoculated with *S. meliloti* Rm1021 despite strong root adhesion in this pairing.

**Figure 1.**
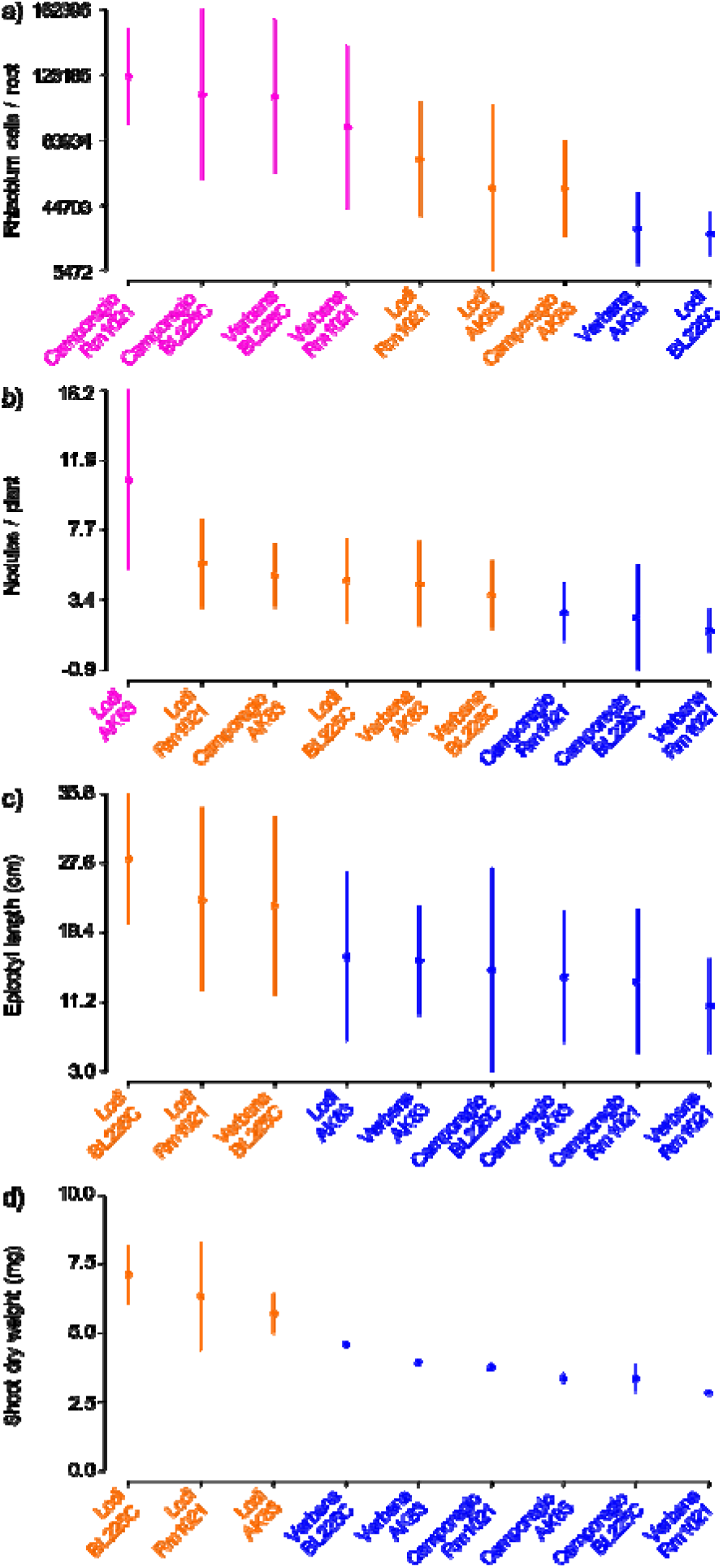
Strain-by-plant variation of symbiosis-associated phenotypes. The number of rhizobium cells retrieved from plant roots (a), number of nodules per plant (b), epicotyl length (c), and the plant dry weight (d), are reported. Different colors (pink, orange, blue) indicate statistically significant groupings (p<0.05) based on a Scott-Knott test. For each condition, the dot indicate the mean value and the vertical lines link the maximum and minimum values.

### Root exudates differ among alfalfa varieties

LC-MS analysis of the alfalfa root exudates detected a total of 2688 unique features, including 392 annotated features, across the two platforms; 1514 hydrophilic features were detected in UPLC-MS positive mode (PP) (288 annotated), and 1174 hydrophilic features were detected in UPLC-MS negative mode (PN) (104 annotated) (**Table S3**). In order to clarify if the metabolite composition of the root exudates from the alfalfa cultivars differed, Principal Component Analysis (PCA) was performed on the two biological replicates of the three cultivars (**Figure 2, Supplemental File S2**). The three cultivars clearly grouped separately from each other, suggesting the presence of a large number of differences in their metabolic compositions. Most of the observed differences were related to amino acids, in particular N-Acetyl-L-leucine, Tryptophan, Cytosine, 3,5-Dihydroxyphenylglycine, and the dipeptide Val-Ala (**Table S4**). Multiple flavones and flavonoids, which include known inducers of NodD activation [66] and of chemotaxis [67], were potentially identified. These include a peak hypothetically attributed to apigenidin (PP_23300) that was found in the Verbena and Camporegio root exudates, liquiritigenin (PP_23583) that was found in the Camporegio and Lodi root exudates, as well as apigenin (PP_25608) and genistein (PP_14051) that were found in variable amounts in the root exudates from all three cultivars. Elemental analysis (CHNS) of root exudates was also performed (**Table S5**), and the results were used to normalize the quantity of root exudate used in the treatment of *S. meliloti* strains, based on equalizing the amount of total organic carbon (TOC) added to each culture.

**Figure 2.**
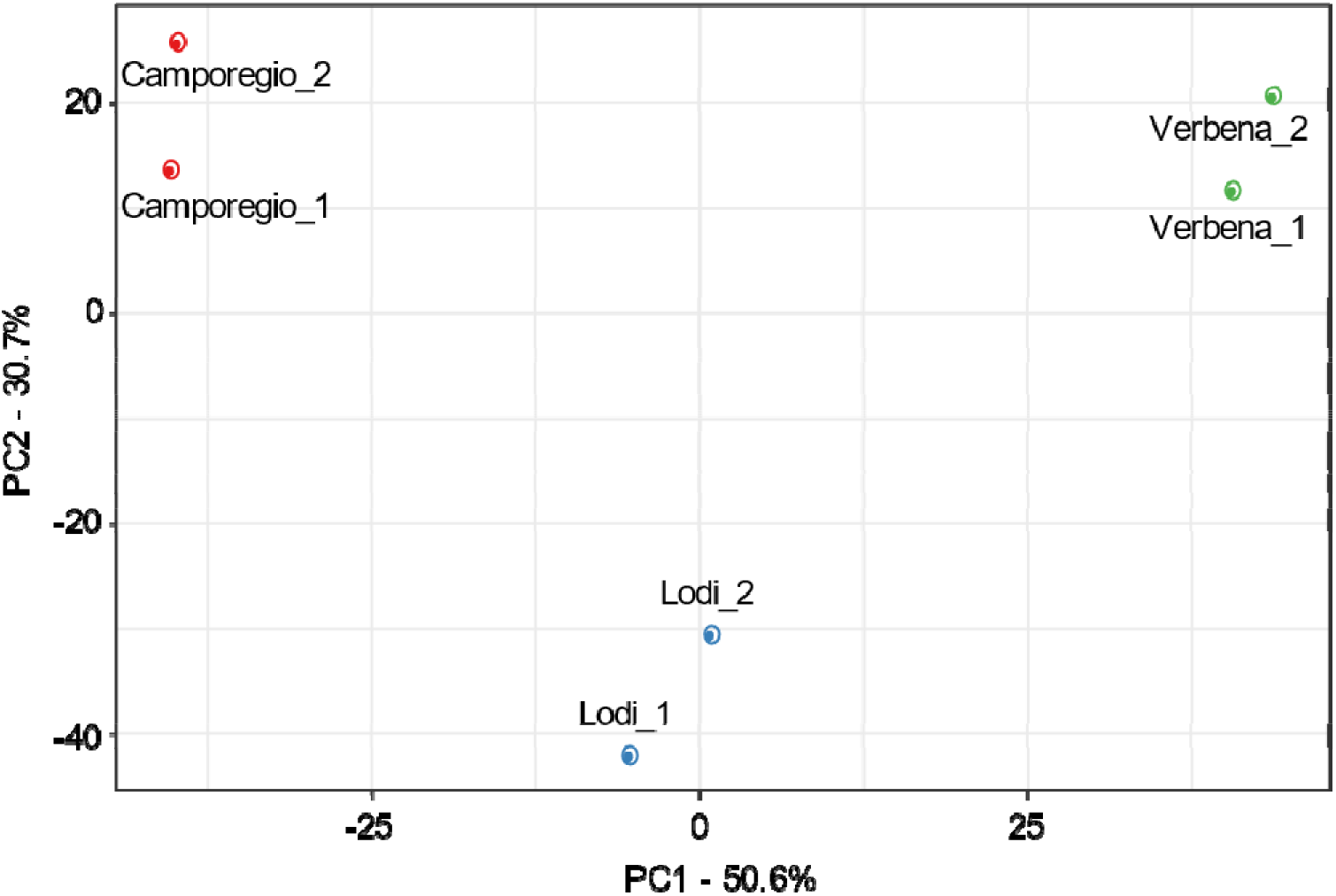
Plant root exudates have different metabolite composition. Principal Component Analysis of 2688 chemical features obtained from LC-MS of root exudates of the Verbena, Lodi, and Camporegio cultivars of alfalfa.

### The number of differentially expressed genes changes in strains x conditions combinations

The global transcriptional responses of the three *S. meliloti* wild type strains following a four-hour exposure to luteolin or alfalfa root exudate was evaluated using RNA-sequencing. In addition, a fourth strain (BM806, referred to as “hybrid” for simplicity) was included [46]; results for this strain will be discussed below. The list of differentially expressed genes (DEGs) for each strain and condition (luteolin and the root exudates of the three plant varieties) against the control (blank sample) is reported in **Table S6**. DEGs were considered to be biologically significant if they had a ≥ 2-fold change in expression and an adjusted *p*-value < 0.01. The numbers of DEGs are shown in **Tables 1** and **S7**. RT-qPCR on a panel of seven DEGs validated the reliability of the RNA-seq data (**Table S8**).

**Table 1.** Significant DEGs. The number of significant DEGs with respect to blank control (2-fold change in expression and an adjusted p-value ≤0.01) and the percentage with respect to the total number of genes is reported.

|  | **Camporegio** | **Lodi** | **Verbena** | **Luteolin** |
| --- | --- | --- | --- | --- |
| **1021** | 516 (8.79%) | 32 (0.55%) | 506 (8.62%) | 36 (0.61%) |
| **AK83** | 357 (5.84%) | 66 (1.08%) | 192 (3.14%) | 60 (0.98%) |
| **BL225C** | 1159 (19.33%) | 76 (1.27%) | 693 (11.56%) | 149 (2.49%) |
| **Hybrid** | 503 (8.38%) | 98 (1.63%) | 325 (5.41%) | 52 (0.87%) |

In general, luteolin treatment resulted in the lowest number of DEGs, ranging from 36 to 149 per strains. Concerning the root exudates, the number of DEGs was influenced by both the strain and the alfalfa cultivar. Overall, the Camporegio and Verbena root exudates induced more gene expression changes than the Lodi root exudate. Cluster analyses of all genes that were differentially expressed in at least one condition (fold change ≥ 2, adjusted *p*-value < 0.01) revealed that, for each strain, the transcriptional responses to the Verbena and Camporegio root exudates were similar, and grouped separately from that of the Lodi cultivar (**Figure 3**; interactive versions are provided in **Supplemental File S1**). Notably, the metabolite composition of the Camporegio and Verbena root exudates were similar along the second principal component (**Figure 2**), suggesting that compounds determining the second principal component of variance may play an important role in modulating *S. meliloti* gene expression. Interestingly, ~ 80% of the genes upregulated by root exudates were found on the chromosomes of the three strains, whereas ~ 77% of the downregulated genes were found on the pSymA and pSymB replicons (**Table S7**). This is consistent with a previous signature-tagged mutagenesis study reporting that 80% of genes required for rhizosphere colonization are chromosomally located in *S. meliloti* Rm1021 [68].

**Figure 3.**
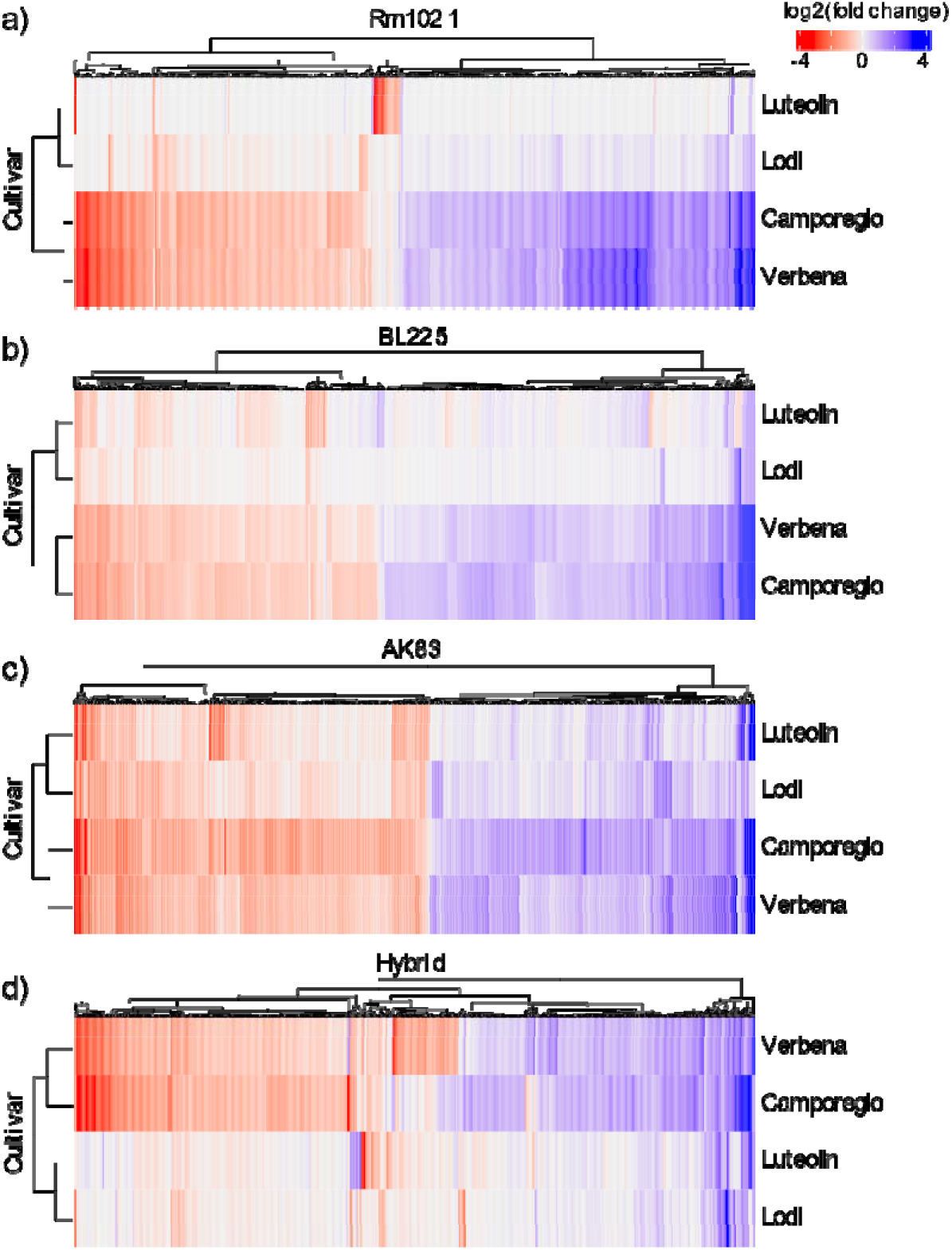
Cluster analysis of the stimulons for the four strains. Differentially expressed genes in each condition are on the columns. a) Rm1021, b) BL225C, c) AK83, d) hybrid strain. See **Supplemental Material file S1** for interactive heatmaps that have each column labelled with the corresponding locus tag

In all conditions, *S. meliloti* BL225C displayed the largest number of DEGs (with up to 20% of genes differentially expressed), while *S. meliloti* AK83 had the fewest. Comparison of the *S. meliloti* Rm1021 data to the NodD3 regulon established elsewhere [69] indicated that 105, 104, and 4 of the DEGs observed in response to the Verbena, Camporegio, and Lodi root exudates, respectively, belong to the NodD3 regulon. However, some of these genes showed contrasting patterns of expression, suggesting the root exudates may also contain antagonistic molecules that repress the *nod* regulon, as previously reported [66, 70]. As the genes overlapping the NodD3 regulon account for ~ 20% or less of the DEGs in each condition, it is likely that most of the observed DEGs belong to *nod*-independent regulons. The majority of DEGs (> 75%) had orthologs in all three of the tested strains (**Table S9**), although expression patterns were not necessarily conserved (**Figure 4**). Interestingly, ≥ 90% of genes upregulated in response to root exudate exposure belonged to the core genome of the three *S. meliloti* strains (**Table S6**), suggesting that the large majority of genes required for alfalfa rhizosphere colonization are highly conserved.

**Figure 4.**
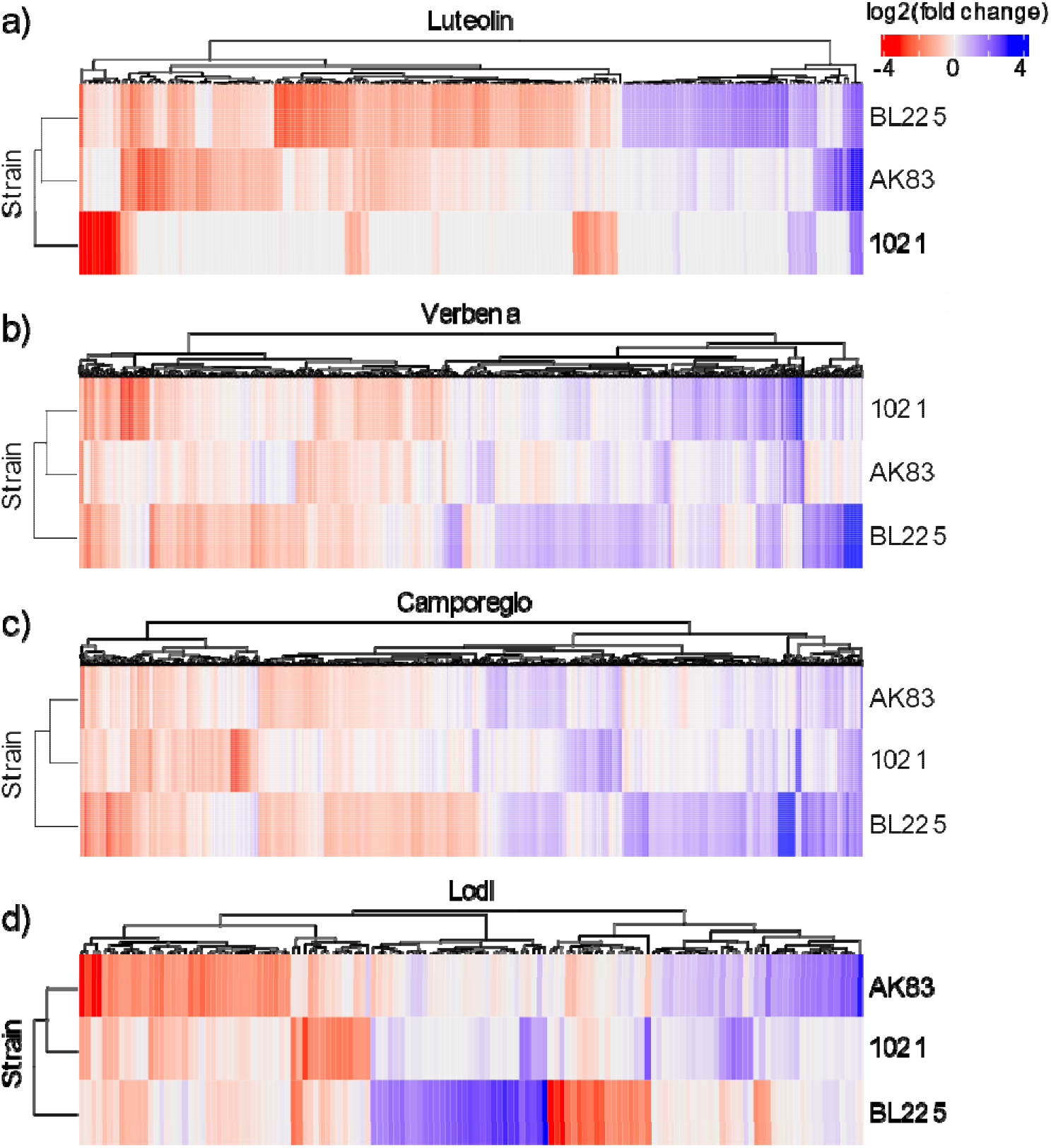
Cluster analysis of the stimulons of the shared set of orthologs. Differentially expressed genes in are on the columns. a) Luteolin, b) Verbena, c) Camporegio, d) Lodi. See **Supplemental Material file S1** for interactive heatmaps that have each column labelled with the corresponding locus tag.

Nested likelihood ratio tests indicated that up to 29% of the conserved genes were influenced by strain x condition interactions, consistent with an important role of GxG interactions in the initiation of rhizobium – legume symbioses (**Table 2**). Moreover, the same analysis emphasized the role of strain genotype in the response to common condition (35% of associated DEGs). Indeed, such strain-by-strain variability on the conserved gene set was also highlighted by the cluster analyses, which indicated that, for each root exudate, the transcriptional responses of *S. meliloti* Rm1021 and AK83 were more similar and grouped separately from that of BL225C (**Figure 4**). Notably, these results do not reflect the genomic relatedness of these strains as Rm1021 and BL225C group together phylogenetically [71].

**Table 2.** Number of expressed genes that showed statistical evidence of each type of expression pattern. Differential expression related to strain, condition, both strain and condition, or interaction between strain and condition is reported. Percentages are calculated on the total number of DEGs. Significance was based on an FDR-corrected p-value < 0.05. * Indicates whether the effect of the tested model on the expression of the gene is significant. A gene can be associated to strain (gene differentially expressed only between strains), condition (gene differentially expressed only between different conditions), strain and condition only (gene differentially expressed in relation to strain and condition but not considering the full model strain x condition), or be associated to the interaction between strain and condition (Strain x condition column). This last situation can be due to significant association to the three tested models ^a^, to the full model and one of the others ^b, c^ or to the full model only ^d^.

|  |  | Groups of DEGs with significant association |  |  |  |  |  |  |  |  |
| --- | --- | --- | --- | --- | --- | --- | --- | --- | --- | --- |
|  |  | Strain | Condi<br>tion | Strain and<br>condition | Strain x<br>condition |  |  |  | None | Total |
| Mod<br>el | Strain | * | - | * | * | * | - | - | - |  |
|  | Condition | - | * | * | * | - | * | - | - |  |
|  | Strain x<br>condition | - | - | - | * | * | * | * | - |  |
| Number of DEGs |  | 2028<br>(25%) | 87<br>(1%) | 1201 (15%) | 180<br>7<sup>a</sup> | 43<br>6<sup>b</sup> | 34<br><sup>c</sup> | 24<sup>d</sup> | 2417<br>(30%) | 8034<br>(100%) |
|  |  |  |  |  | 2301 (29%) |  |  |  |  |  |

**Table 3.** Selected COG categories over- or under-represented among the DEGs. Values represent the log2 fold change in abundance of genes annotated with the given COG category relative to the amount expected by chance. Dashes indicate that the COG category is not statistically different than chance in the given condition (significance threshold: *p*-value ≤ 0.05).

| COG Category | Luteolin |  |  | Camporegio |  |  | Verbena |  |  | Lodi |  |  |
| --- | --- | --- | --- | --- | --- | --- | --- | --- | --- | --- | --- | --- |
|  | Rm1021 | BL225C | AK83 | Rm1021 | BL225C | AK83 | Rm1021 | BL225C | AK83 | Rm1021 | BL225C | AK83 |
| Upregulated genes |  |  |  |  |  |  |  |  |  |  |  |  |
| G | - | - | - | -1.14 | -1.59 | -1.67 | -1.30 | -1.64 | - | - | - | - |
| J | - | - | - | 2.81 | 0.92 | 2.50 | 2.80 | 1.16 | 1.81 | - | - | - |
| N | - | - | - | 4.19 | - | - | 4.25 | - | - | - | - | - |
| O | - | - | 2.38 | 1.56 | 1.44 | 1.50 | 1.44 | 1.91 | 2.21 | - | - | 2.96 |
| Downregulated genes |  |  |  |  |  |  |  |  |  |  |  |  |
| C | - | - | - | 1.88 | 0.88 | 1.74 | 1.89 | 1.11 | 1.79 | 2.71 | - | - |

### Stimulons differ in the set of elicited functions

Functional enrichment analyses, based on KEGG modules and COG categories, were performed to give a global overview of the functions of the DEGs (**Supplemental File S2**). Strain and condition specific patterns of functional enrichment were observed, consistent with a functional differentiation of the stimulons of each experiment. Nevertheless, a core set of COG categories were commonly over- or under-represented in all three *S. meliloti* strains during exposure to the Camporegio or Verbena root exudates. These included an enrichment among the up-regulated genes of COG categories J and O related to protein expression and modification, suggesting that the root exudates stimulated a major remodelling of the proteome. In addition, the COG category G (carbohydrate transport and metabolism) was under-represented among the up-regulated genes while the COG category C (energy production and conversion) was over-represented among the down-regulated genes. This observation suggests that the root exudates stimulated a global change in the cellular energy production pathways versus growth in our standard minimal medium with succinate as the sole carbon source.

Among the mostly highly expressed genes in *S. meliloti* Rm1021 during exposure to the Verbena and Camporegio root exudates were *smc03024* and *smc03028*, encoding components of the flagellar apparatus (*flgF* and *flgC*, respectively); the orthologs of these genes were not induced in BL225C nor AK83. The induction of motility is in contrast the observation that luteolin alone decreases the motility of *S. meliloti* Rm1021 strain [69, 72]. Presumably, this reflects the presence of additional stimuli in the root exudates. Indeed, amino acids present in root exudates are known to stimulate chemiotactic behaviour in *S. meliloti* [73] and a signature tagged mutagenesis showed that motility-related genes are relevant during competition for rhizosphere colonization in *S. meliloti* Rm1021 [68].

Differences in the transcriptomes of two *Bradyrhizobium diazoefficiens* strains exposed to root exudates were suggested to be related to differences in their competitive abilities [74]. We therefore examined the expression patterns of several genes likely to play a role in competition for rhizosphere colonization and root adhesion. It was previously suggested that the *sin* quorum sensing system is involved in competition in *S. meliloti* [75]; in our data, *sinI* (*smc00168*) was repressed in *S. meliloti* Rm1021 in the presence of the Camporegio and Verbena root exudates, but no changes in the expression of the orthologous genes in strains AK83 or BL225C were observed. No evidence was found in any of the strains for changes in expression of galactoglucan or succinoglucan biosynthesis genes, such as *wgaA* (*sm_b21319)* and *wgeA* (*sm_b21314*). The Verbena and Camporegio root exudates induced expression of the rhizobactin transport gene (*sma2337* [*rhtX*]) of Rm1021 and BL225C; this gene is not found in AK83. This may be a consequence of the root exudates chelating the available iron [76], consequently eliciting siderophore production that can inhibit growth of strains lacking siderophores [77]. Plasmid pSINME01 of *S. meliloti* AK83 exhibits similarity with the plasmid pHRC017 from *S. meliloti* C017, which confers a competitive advantage for nodule occupancy and host range restrictions [78]. Considering that a few of the genes on the plasmids pSINME01 and pSINME02 were differentially expressed upon exposure to root exudates, it is possible that the accessory plasmids of strain AK83 also contribute to competition for rhizosphere colonization [78].

Differences in gene expression patterns across conditions may be related, in part, to differences in the presence of flavonoids. For example, the *emrAB* systems (*smc03167* and *smc03168*) is known to be induced by luteolin and apigenin [79]. Here, these genes were induced by luteolin and the Camporegio and Verbena root extracts that putatively contained apigenin, but they were not induced by the Lodi root extract that lacked apigenin.

### Mobilization of the symbiotic megaplasmid results in nonadditive changes in stimulons

To evaluate the impact of inter-replicon epistatic interactions on the transcriptional response of *S. meliloti* to alfalfa root exudates, we used RNA-seq to characterize the response of a *S. meliloti* hybrid strain, containing the chromosome and pSymB of strain Rm2011 and the symbiotic megaplasmid (pSINMEB01) of strain BL225C [46]. Cluster analyses clearly demonstrated that the transcriptome of the hybrid strain differed from both the BL225C and Rm1021 wild type strains in all conditions (**Figure 5**). Of particular interest were the results observed during exposure to the Lodi root exudate. We previously showed that alfalfa cv. Lodi plants inoculated with the hybrid strain were larger than those inoculated with either BL225C or Rm1021 [44]. Here, we observed that exposure to Lodi root exudate results in more differentially expressed genes in the hybrid strain (98 genes) than in either Rm1021 or BL225C (32 and 76 genes, respectively; **Table 1**). In particular, a cluster of genes was specifically upregulated in the hybrid strain, and the majority of these genes were located on the symbiotic megaplasmid. The presence of these nonadditive transcriptional changes may reflect a loss of regulation of these megaplasmid genes by chromosomal regulators [80, 81], providing a potential molecular mechanism underlying the improved symbiotic phenotype of the hybrid compared to both wild type strains.

**Figure 5.**
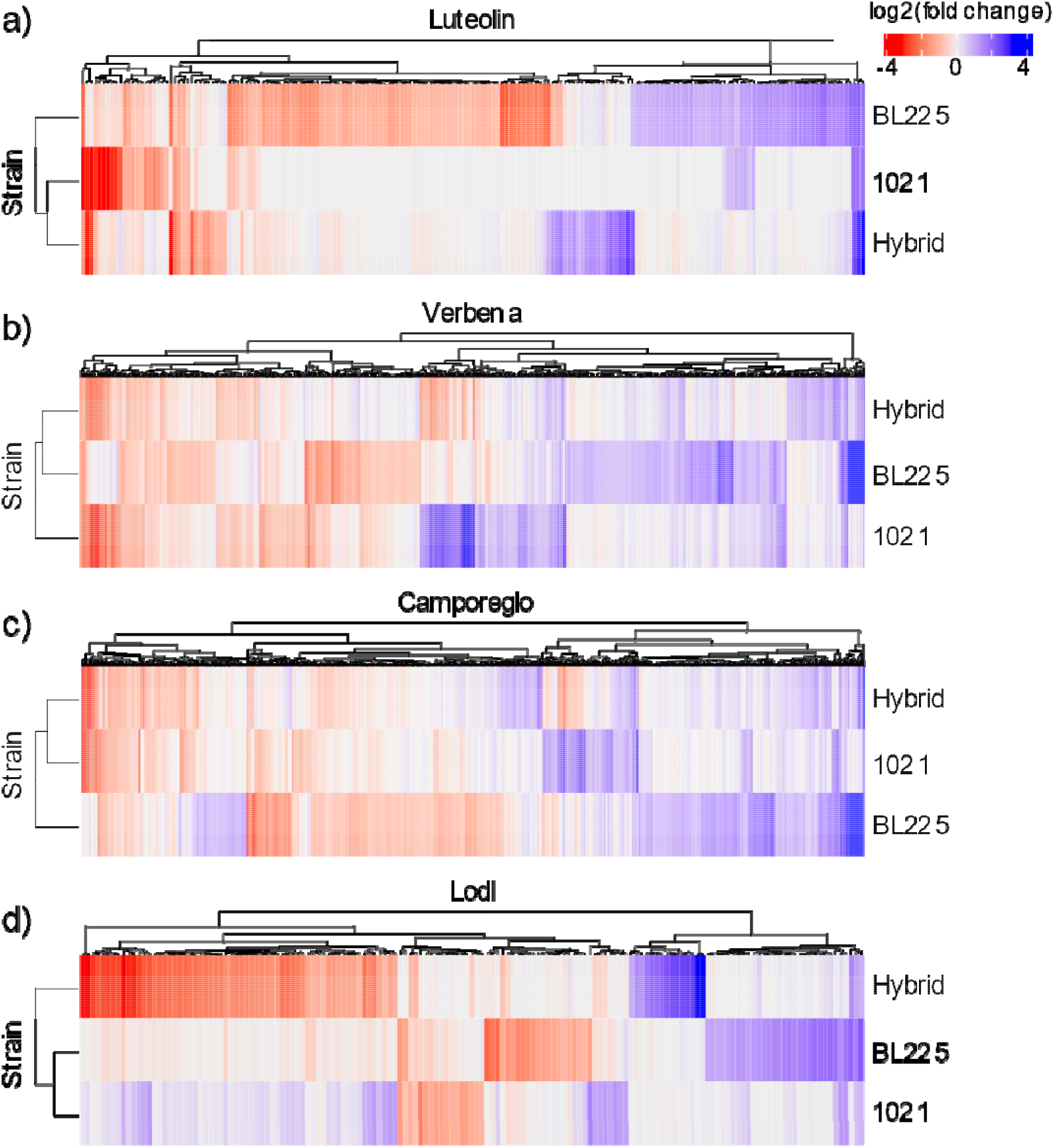
Clustering of the pSymA-hybrid strain with respect to the two parental ones (Rm1021 and BL225C). Heatmaps based on DEGs of orthologous genes. a) Luteolin, b) Verbena, c) Camporegio, d) Lodi. See **Supplemental Material file S1** for interactive heatmaps that have each column labelled with the corresponding locus tag.

## Discussion

Rhizobium-legume interactions are complex multistep phenomena, that begin with an exchange of signals between two partners [16, 82]. The rhizobia initially detect the plant through perception of flavonoids in the root exudate of legumes by NodD proteins, which then triggers the production of lipochitooligosaccharide molecules known as Nod factors. Nod factors are then recognized by specific LysM receptor kinase proteins in plant root cells, triggering the symbiosis signalling pathway and initiating the formation of a nodule. However, root exudates contain a mixture of flavonoids, some of them having different agonistic activity on NodD [66]. Root exudates also contain many other molecules that can serve as signals or support rhizobium metabolism, such as amino acids and sugars, that may influence the ability of rhizobia to successfully colonize the rhizosphere and be in a position to enter into the symbiosis [83, 84]. Consequently, interactions between plant and rhizobium genotypes are expected to influence the success of the initial interaction between the two partners.

Previous works have identified a clear role for GxG interactions in the partnership between *S. meliloti* and *M. trucantula* [85], demonstrating that aerial biomass was influenced by the plant and rhizobium genotypes as well as by their interaction. Here, we demonstrated that GxG interactions also have a significant impact on the adherence of *S. meliloti* strains to alfalfa roots, as a representative phenotype for an early stage of the interaction between these partners. Rhizosphere colonization appears to have a direct impact on nodule colonization [68, 86]; while our data does not address if root adhesion is correlated with competition for nodule occupancy in mixed inoculums, it does suggest that root adhesion is poorly correlated with overall symbiotic efficiency in single-inoculum studies. Previous studies have also demonstrated an influence of GxG interactions on the nodule transcriptome of *Medicago* – *Sinorhizobium* symbioses [33, 34]. Here, we showed that GxG interactions similarly have an important contribution in determining the transcriptional response of *S. meliloti* to detection of *M. sativa* root exudates. Together, these results demonstrate that GxG interactions have a meaningful impact on the outcome of rhizobium – legume symbioses at multiple stages of development.

Exposure of *B. diazoefficiens* to soybean root exudates resulted in changes in expression of 450 genes, representing nearly 5.6% of the genome, and the impact of soybean root exudate differed between the two tested *B. diazoefficiens* strains [74]. Similarly, between 0.5% and 20% of *S. meliloti* genes were differentially expressed following exposure to alfalfa root exudate, depending on the host – symbiont combination. The similarities/differences in response of the three *S. meliloti* strains to treatments did not appear to depend the phylogenetic relatedness of the strains [71], although this cannot be definitively concluded without analysis of additional strains. Nevertheless, these results emphasize the importance of transcriptional rewiring during strain diversification in bacteria [81]. Similarly, studies with eukaryotic organisms indicate that adaptation has an important role in differentiating the gene expression patterns of organisms [87, 88].

The root exudate stimulons only partially overlapped with the stimulons of luteolin, a known inducer of NodD in *S. meliloti* [66], confirming that alfalfa root exudates contain numerous molecular signals aside form flavonoids that may influence the competitiveness of various rhizobium strains. Importantly, the transcriptional patterns induced by alfalfa root exudates differed depending on the cultivar from which they were collected; whether these differences are adaptive requires further investigation. Additionally, there was a clear relationship between the differences in the *S. meliloti* gene expression profiles and the overall chemical similarity of the root exudates as measured with LC-MS; the Camporegio and Verbena root exudates induced similar gene expression changes, while also being similar along the second principal component of variance (accounting for 30% of the variance) in the PCA of the root exudate composition. In future work, it would be interesting to define which compounds in the root exudates have the greatest impact on the *S. meliloti* transcriptome.

In addition to the impact of GxG interactions on rhizobium – legume symbioses, there is a potential for inter-replicon interactions within rhizobium genomes to further influence the symbiosis. Indeed, inter-replicon epistatic interactions are abundant in the *S. meliloti* genome [89]. To address the contribution of inter-replicon interactions on symbiosis, we examined a hybrid strain in which the symbiotic megaplasmid of *S. meliloti* Rm2011 (a near identical strain to Rm1021 [43]) was replaced with the symbiotic megaplasmid of *S. meliloti* BL225C. Non-additive effects on the transcriptional profiles associated with all three replicons were observed in the hybrid strain relative to Rm1021 and BL225C, indicating that megaplasmid mobilization induced a global rewiring of gene expression likely due to transcriptional cross-talk among the replicons [81, 90]. Similarly, non-additive effects on the transcriptome of plant hybrids have been extensively explored [91] and demonstrated as one of the basis for heterosis in crops [92]. The results with the hybrid lead us to hypothesize that the large symbiotic variability observed in natural *S. meliloti* isolates may partly be related to genome-wide transcriptome changes following large-scale horizontal gene transfer followed by natural selection. If true, however, this would limit our ability to predict the competitiveness of rhizobium isolates from their genome sequence.

In conclusion, this study demonstrated that the initial perception of legumes by rhizobia leads to hundreds of changes in the rhizobium transcriptome, and that these changes are dependent on the plant genotype, the rhizobium genotype, and genotype x genotype interactions. These results complement past studies demonstrating a role of GxG interactions in determining the transcriptome of both the legume and rhizobium partners in mature N_2_-fixing nodules [33, 34]. The majority of genes up-regulated in response to alfalfa root exudates were conserved in all three strains, supporting the hypothesis that the *S. meliloti* lineage was adapted to rhizosphere colonization before gaining the genes required for symbiotic nitrogen fixation [68]. Additionally, the transcriptional response to perception of alfalfa root exudate involved genes from all three of the *S. meliloti* replicons, and seemingly involved non-additive effects resulting from inter-replicon interactions.

## Acknowledgments

We are grateful to Gabriele Brazzini for technical assistance in setting up the root adhesion test. This work was supported by Fondazione Cassa di Risparmio di Firenze, grant n. 18204, 2017.0719, by “MICRO4Legumes” grant (Italian Ministry of Agriculture) and by the grant as “Dipartimento di Eccellenza 2018-2022” by the Italian Ministry of Education, University and Research (MIUR). LC was supported by a fellowship from MICRO4Legumes (Italian Ministry of Agriculture). Work in the GCD laboratory is supported by funding from Queen’s University and the Natural Sciences and Engineering Research Council of Canada.

## Supplementary Material

**Supplemental Material file S1**. Interactive heatmaps of the stimulons. A .zip folder containing .html files for detailed descriptions of Figures 3, 4 and 6.

**Supplemental Material file S2**. Functional categorization of DEGs. An Excel file containing four datasheets with KEGG and COG categories found in up and downregulated genes.

**Supplemental Material file S3**. A .zip archive containing the following tables and figures (see below)

## Supplementary tables

**Table S1**. List of strains and alfalfa cultivars. .docx file.

**Table S2**. Primers used in this study. .docx file.

**Table S3**. List of LC-MS peaks. Excel file.

**Table S4**. Metabolites with the greatest differences among root exudates. Results of Simper analysis based on the decomposition of the Bray-Curtis dissimilarity obtained from each peak ID value .docx file.

**Table S5**. Chemical composition of root exudates from CHN analysis. .docx file.

**Table S6**. Significant differentially expressed genes (DEGs). .csv file.

**Table S7**. Overall number of DEGs in the strains by treatment combinations, their location on the *S. meliloti* replicons and up and downregulation with respect to blank control. Excel file.

**Table S8**. Results of quantitative RT-PCR on selected genes. .docx file.

**Table S9**. Pangenome ortholog assignement from Roary. .csv file.

## Supplementary Figures

**Figure S1**. Symbiosis-associated phenotypes. The number of rhizobium cells retrieved from plant roots (a), number of nodules per plant (b), epicotyl length (c), and the plant dry weight (d), are reported. Letters indicate groupings based on Tukey contrasts (p<0.05). Error bars indicate one standard deviation. .png file

**Figure S2**. Plot of Principal Component Analysis from LC-MS of root exudates of the Verbena, Lodi and Camporegio cultivars of alfalfa, including the blank control. .png file

## References

1. Rosenberg E, Zilber-Rosenberg I. Microbes Drive Evolution of Animals and Plants: the Hologenome Concept. MBio 2016; 7: e01395–15-.

2. Mus F, Crook MB, Garcia K, Costas AG, Geddes BA, Kouri ED, et al. Symbiotic nitrogen fixation and the challenges to its extension to nonlegumes. Appl Environ Microbiol . 2016., 82

3. Lee YK, Mazmanian SK. Has the Microbiota Played a Critical Role in the Evolution of the Adaptive Immune System? Science (80-) 2010; 330: 1768 LP – 1773.

4. Jones BW, Nishiguchi MK. Counterillumination in the Hawaiian bobtail squid, Euprymna scolopes Berry (Mollusca: Cephalopoda). Mar Biol 2004; 144: 1151–1155.

5. Nyholm S V, McFall-Ngai M. The winnowing: establishing the squid–vibrio symbiosis. Nat Rev Microbiol 2004; 2: 632–642.

6. Kiers ET, Rousseau RA, West SA, Denison RF. Host sanctions and the legume – rhizobium mutualism. Nature 2003; 425: 1095–1098.

7. Mwafulirwa L, Baggs EM, Russell J, George T, Morley N, Sim A, et al. Barley genotype influences stabilization of rhizodeposition-derived C and soil organic matter mineralization. Soil Biol Biochem 2016; 95: 60–69.

8. Escudero-Martinez C, Bulgarelli D. Tracing the evolutionary routes of plant–microbiota interactions. Curr Opin Microbiol 2019; 49: 34–40.

9. Pasolli E, Asnicar F, Manara S, Quince C, Huttenhower C, Correspondence NS, et al. Extensive Unexplored Human Microbiome Diversity Revealed by Over 150,000 Genomes from Metagenomes Spanning Age, Geography, and Lifestyle. Cell 2019; 176: 1–14.

10. Singh BK, Liu H. Eco-holobiont◻: A new concept to identify drivers of host-associated microorganisms. 2019; 00.

11. de Souza RSC, Armanhi JSL, Damasceno N de B, Imperial J, Arruda P. Genome Sequences of a Plant Beneficial Synthetic Bacterial Community Reveal Genetic Features for Successful Plant Colonization. Front Microbiol 2019; 10.

12. Levy A, Gonzalez IS, Mittelviefhaus M, Clingenpeel S, Paredes SH, Miao J, et al. Genomic features of bacterial adaptation to plants. Nat Genet 2018; 50: 138–150.

13. Pini F, Galardini M, Bazzicalupo M, Mengoni A. Plant-Bacteria Association and Symbiosis: Are There Common Genomic Traits in Alphaproteobacteria? Genes (Basel) 2011; 2: 1017–1032.

14. Guttman D, McHardy AC, Schulze-Lefert P. Microbial genome-enabled insights into plant– microorganism interactions. Nat Rev Genet 2014; 15: 797–813.

15. Poole P, Ramachandran V, Terpolilli J. Rhizobia: from saprophytes to endosymbionts. Nat Publ Gr 2018.

16. Oldroyd GED. Speak, friend, and enter: signalling systems that promote beneficial symbiotic associations in plants. Nat Rev Microbiol 2013; 11: 252–63.

17. Gage DJ. Infection and Invasion of Roots by Symbiotic, Nitrogen-Fixing Rhizobia during Nodulation of Temperate Legumes. Society 2004; 68: 280–300.

18. Udvardi M, Poole PS. Transport and metabolism in legume-rhizobia symbioses. Annu Rev Plant Biol 2013; 64: 781–805.

19. Kereszt A, Mergaert P, Kondorosi E. Bacteroid development in legume nodules: Evolution of mutual benefit or of sacrificial victims? Mol Plant-Microbe Interact 2011; 24: 1300–1309.

20. Benezech C, Doudement M, Gourion B. Legumes tolerance to rhizobia is not always observed and not always deserved. Cell Microbiol 2019; 1–9.

21. Gibson KE, Kobayashi H, Walker GC. Molecular Determinants of a Symbiotic Chronic Infection. Annu Rev Genet 2008; 42: 413–441.

22. Burghardt LT. Evolving together, evolving apart: measuring the fitness of rhizobial bacteria in and out of symbiosis with leguminous plants. New Phytol 2019.

23. Paffetti D, Scotti C, Gnocchi S, Fancelli S. Genetic Diversity of an Italian Rhizobium meliloti Population from Different Medicago sativa Varieties. Appl Environ Microbiol 1996; 62: 2279–2285.

24. Carelli M, Gnocchi S, Fancelli S, Mengoni A, Paffetti D, Scotti C, et al. Genetic diversity and dynamics of Sinorhizobium meliloti populations nodulating different alfalfa cultivars in Italian soils. Appl Environ Microbiol 2000; 66.

25. Rangin C, Brunel B, Perrineau M, Béna G. Effects of Medicago truncatula genetic diversity, rhizobial competition and strain effectiveness on the diversity of a natural Sinorhizobium spp. community. Appl Environ Microbiol Microbiol 2008; 74: 5653–5661.

26. Heath KD, Tiffin P. Context dependence in the coevolution of plant and rhizobial mutualists. Proc R Soc B Biol Sci 2007; 274: 1905–1912.

27. Heath KD, Tiffin P. Stabilizing mechanisms in a legume-rhizobium mutualism. Evolution (N Y) 2009; 63: 652–662.

28. Heath KD. Intergenomic epistasis and coevolutionary constraint in plants and rhizobia. Evolution (N Y) 2010; 64: 1446–1458.

29. Ehinger M, Mohr TJ, Starcevich JB, Sachs JL, Porter SS, Simms EL. Specialization-generalization trade-off in a Bradyrhizobium symbiosis with wild legume hosts. BMC Ecol 2014; 14.

30. Burghardt LT, Epstein B, Guhlin J, Nelson MS, Taylor MR, Young ND, et al. Select and resequence reveals relative fitness of bacteria in symbiotic and free-living environments. Proc Natl Acad Sci 2018; 115: 2425–2430.

31. Burghardt LT, Trujillo DI, Epstein B, Tiffin P, Young ND. A Select and Resequence Approach Reveals Strain-Specific Effects of Medicago Nodule-Specific PLAT-Domain Genes. Plant Physiol 2020; 182: 463–471.

32. Roux B, Rodde N, Jardinaud MF, Timmers T, Sauviac L, Cottret L, et al. An integrated analysis of plant and bacterial gene expression in symbiotic root nodules using laser-capture microdissection coupled to RNA sequencing. Plant J 2014; 77: 817–837.

33. Heath KD, Burke PV, Stinchombe JR. Coevolutionary genetic variation in the legume-rhizobium transcriptome. Mol Ecol 2012; 19: 4735–4747.

34. Burghardt LT, Guhlin J, Chun CL, Liu J, Sadowsky MJ, Stupar RM, et al. Transcriptomic basis of genome by genome variation in a legume-rhizobia mutualism. Mol Ecol 2017; 26: 6122–6135.

35. Triplett EW, Sadowsky MJ. Genetics of competition for nodulation of legumes. Annu Rev Microbiol 1992; 46: 399–422.

36. Checcucci A, DiCenzo GC, Bazzicalupo M, Mengoni A. Trade, diplomacy, and warfare: The Quest for elite rhizobia inoculant strains. Front Microbiol 2017; 8.

37. Sprent JI, Ardley J, James EK. Biogeography of nodulated legumes and their nitrogen-fixing symbionts. New Phytol 2017; 215: 40–56.

38. Frame J, Charlton JFL, Laidlaw AS. Temperate forage legumes. 1998. Cab International, Wallingford, UK.

39. Galibert F, Finan TM, Long SR, Puhler A, Abola P, Ampe F, et al. The composite genome of the legume symbiont Sinorhizobium meliloti. Science (80-) 2001; 293: 668–672.

40. Harrison PW, Lower RPJ, Kim NKD, Young JPW. Introducing the bacterial ‘chromid’: Not a chromosome, not a plasmid. Trends Microbiol 2010; 18: 141–148.

41. Meade HM, Long SR, Ruvkun GB, Brown SE, Ausubel FM. Physical and genetic characterization of symbiotic and auxotrophic mutants of Rhizobium meliloti induced by transposon Tn5 mutagenesis. J Bacteriol 1982; 149: 114 LP – 122.

42. Galardini M, Mengoni A, Brilli M, Pini F, Fioravanti A, Lucas S, et al. Exploring the symbiotic pangenome of the nitrogen-fixing bacterium Sinorhizobium meliloti. BMC Genomics 2011; 12: 235.

43. Meade HM, Signer ER. Genetic mapping of Rhizobium meliloti. Proc Natl Acad Sci 1977; 74: 2076–2078.

44. Checcucci A, Dicenzo GC, Ghini V, Bazzicalupo M, Becker A, Decorosi F, et al. Creation and Characterization of a Genomically Hybrid Strain in the Nitrogen-Fixing Symbiotic Bacterium Sinorhizobium meliloti. ACS Synth Biol 2018; 7: 2365–2378.

45. Teuber LR, Taggard KL, Gibbs LK, McCaslin MH, Peterson MA, Barnes DK. Fall dormancy. Stand. tests to Charact. alfalfa Cultiv. CC Fox) p. A-1.(North Am. Alfalfa Improv. Conf. Beltsville, MD). 1998.

46. Checcucci A, diCenzo G, Ghini V, Bazzicalupo M, Becker A, Decorosi F, et al. Creation and characterization of a genomically hybrid strain in the nitrogen-fixing symbiotic bacterium Sinorhizobium meliloti. ACS Synth Biol 2018.

47. Trabelsi D, Pini F, Aouani ME, Bazzicalupo M, Mengoni A. Development of real-time PCR assay for detection and quantification of Sinorhizobium meliloti in soil and plant tissue. Lett Appl Microbiol 2009; 48: 355–361.

48. Checcucci A, Azzarello E, Bazzicalupo M, Galardini M, Lagomarsino A, Mancuso S, et al. Mixed nodule infection in Sinorhizobium meliloti-medicago sativa symbiosis suggest the presence of cheating behavior. Front Plant Sci 2016; 7.

49. R Development Core Team. R: A language and environment for statistical computing. R Foundation for Statistical Computing, Vienna, Austria. ISBN 3-900051-07-0, URL http://www.R-project.org/. R Found Stat Comput Vienna, Austria 2012.

50. Checcucci A, Azzarello E, Bazzicalupo M, Carlo AD, Emiliani G, Mancuso S, et al. Role and regulation of ACC deaminase gene in Sinorhizobium melilotr: Is it a symbiotic, rhizospheric or endophytic gene? Front Genet 2017; 8.

51. Giavalisco P, Ko□hl K, Hummel J, Seiwert B, Willmitzer L. 13C isotope-labeled metabolomes allowing for improved compound annotation and relative quantification in liquid chromatography-mass spectrometry-based metabolomic research. Anal Chem 2009; 81: 6546–6551.

52. Oksanen J, Blanchet F, Kindt R, Legendre P, Minchin P, O’Hara R, et al. vegan: Community Ecology Package. R package version 2.0-10. R Packag version . 2013., 1: 10.4135/9781412971874.n145

53. Bacci G, Bazzicalupo M, Benedetti A, Mengoni A. StreamingTrim 1.0: A Java software for dynamic trimming of 16S rRNA sequence data from metagenetic studies. Mol Ecol Resour 2014; 14.

54. Patro R, Duggal G, Love MI, Irizarry RA, Kingsford C. Salmon provides fast and bias-aware quantification of transcript expression. Nat Methods 2017; 14: 417–419.

55. Soneson C, Love MI, Robinson MD. Differential analyses for RNA-seq: transcript-level estimates improve gene-level inferences. F1000Research 2015; 4.

56. Love MI, Huber W, Anders S. Moderated estimation of fold change and dispersion for RNA-seq data with DESeq2. Genome Biol 2014; 15: 550.

57. Page AJ, Cummins CA, Hunt M, Wong VK, Reuter S, Holden MTG, et al. Roary: Rapid large-scale prokaryote pan genome analysis. Bioinformatics 2015; 31: 3691–3693.

58. Gu Z, Eils R, Schlesner M. Complex heatmaps reveal patterns and correlations in multidimensional genomic data. Bioinformatics 2016; 32: 2847–2849.

59. Galili T, O’Callaghan A, Sidi J, Sievert C. heatmaply: an R package for creating interactive cluster heatmaps for online publishing. Bioinformatics 2018; 34: 1600–1602.

60. Wickham Hadley. ggplot2: Elegant Graphics for Data Analysis. 2009. Springer-Verlag New York.

61. Huerta-Cepas J, Szklarczyk D, Heller D, Hernández-Plaza A, Forslund SK, Cook H, et al. eggNOG 5.0: a hierarchical, functionally and phylogenetically annotated orthology resource based on 5090 organisms and 2502 viruses. Nucleic Acids Res 2019; 47: D309–D314.

62. Huerta-Cepas J, Forslund K, Coelho LP, Szklarczyk D, Jensen LJ, Von Mering C, et al. Fast genome-wide functional annotation through orthology assignment by eggNOG-mapper. Mol Biol Evol 2017; 34: 2115–2122.

63. Reback J, McKinney W, jbrockmendel, Bossche J Van den, Augspurger T, Cloud P, et al. pandas-dev/pandas: Pandas 1.0.3. 2020.

64. Oliphant TE. A guide to NumPy. 2006. Trelgol Publishing USA.

65. Biondi EG, Tatti E, Comparini D, Giuntini E, Mocali S, Giovannetti L, et al. Metabolic Capacity of Sinorhizobium (Ensifer) meliloti Strains as Determined by Phenotype MicroArray Analysis L †. Society 2009; 75: 5396–5404.

66. Peck MC, Fisher RF, Long SR. Diverse Flavonoids Stimulate NodD1 Binding to nod Gene Promoters in Sinorhizobium meliloti. J Bacteriol 2006; 188: 5417–5427.

67. Caetano-Anollés G, Crist-Estes DK, Bauer WD. Chemotaxis of Rhizobium meliloti to the plant flavone luteolin requires functional nodulation genes. J Bacteriol 1988; 170: 3164 LP – 3169.

68. Salas ME, Lozano MJ, López JL, Draghi WO, Serrania J, Torres Tejerizo GA, et al. Specificity traits consistent with legume-rhizobia coevolution displayed by Ensifer meliloti rhizosphere colonization. Environ Microbiol 2017; 19: 3423–3438.

69. Barnett MJ, Long SR. The Sinorhizobium meliloti SyrM regulon: effects on global gene expression are mediated by syrA and nodD3. J Bacteriol 2015; 197: 1792–1806.

70. Zuanazzi JAS, Clergeot PH, Quirion J-C, Husson H-P, Kondorosi A, Ratet P. Production of Sinorhizobium meliloti nod gene activator and repressor flavonoids from Medicago sativa roots. Mol plant-microbe Interact 1998; 11: 784–794.

71. Galardini M, Pini F, Bazzicalupo M, Biondi EG, Mengoni A. Replicon-dependent bacterial genome evolution: The case of Sinorhizobium meliloti. Genome Biol Evol 2013; 5: 542–558.

72. Spini G, Decorosi F, Cerboneschi M, Tegli S, Mengoni A, Viti C, et al. Effect of the plant flavonoid luteolin on Ensifer meliloti 3001 phenotypic responses. Plant Soil 2016; 399.

73. Webb BA, Helm RF, Scharf BE. Contribution of individual chemoreceptors to Sinorhizobium meliloti chemotaxis towards amino acids of host and nonhost seed exudates. Mol Plant-Microbe Interact 2016; 29: 231–239.

74. Liu Y, Jiang X, Guan D, Zhou W, Ma M, Zhao B, et al. Transcriptional analysis of genes involved in competitive nodulation in Bradyrhizobium diazoefficiens at the presence of soybean root exudates. Sci Rep 2017; 7: 10946.

75. McIntosh M, Krol E, Becker A. Competitive and cooperative effects in quorum-sensing-regulated galactoglucan biosynthesis in Sinorhizobium meliloti. J Bacteriol 2008; 190: 5308–5317.

76. Parker DR, Reichman SM, Crowley DE. Metal Chelation in the Rhizosphere. Roots Soil Manag Interact between Roots Soil . 2005., 57–93

77. diCenzo GC, MacLean AM, Milunovic B, Golding GB, Finan TM. Examination of Prokaryotic Multipartite Genome Evolution through Experimental Genome Reduction. PLoS Genet 2014; 10.

78. Crook MB, Lindsay DP, Biggs MB, Bentley JS, Price JC, Clement SC, et al. Rhizobial Plasmids That Cause Impaired Symbiotic Nitrogen Fixation and Enhanced Host Invasion. Mol Plant-Microbe Interact 2012; 25: 1026–1033.

79. Rossbach S, Kunze K, Albert S, Zehner S, Göttfert M. The Sinorhizobium meliloti EmrAB efflux system is regulated by flavonoids through a TetR-like regulator (EmrR). Mol Plant-Microbe Interact 2014; 27: 379–387.

80. Ramos JL, Marqués S, Timmis KN. Transcriptipnal control of the Pseudomonas TOL plasmid catabolic operons is achieved through an interplay of host factors and plasmid-encoded regulators. Annu Rev Microbiol 1997; 51: 341–373.

81. Galardini M, Brilli M, Spini G, Rossi M, Roncaglia B, Bani A, et al. Evolution of Intra-specific Regulatory Networks in a Multipartite Bacterial Genome. PLoS Comput Biol 2015; 11: e1004478.

82. Oldroyd GED, Murray JD, Poole PS, Downie JA. The Rules of Engagement in the Legume-Rhizobial Symbiosis. Annu Rev Genet 2011; 45: 119–144.

83. Barbour WM, Hattermann DR, Stacey G. Chemotaxis of Bradyrhizobium japonicum to soybean exudates. Appl Environ Microbiol 1991; 57: 2635–2639.

84. Webb BA, Compton KK, Martin JS, Taylor D, Sobrado P, Scharf BE. Sinorhizobium meliloti Chemotaxis to Multiple Amino Acids Is Mediated by the Chemoreceptor McpU. Mol Plant Microbe Interact 2017; 30: 770–777.

85. Mhadhbi H, Jebara M, Limam F, Huguet T, Aouani ME. Interaction between Medicago truncatula lines and Sinorhizobium meliloti strains for symbiotic efficiency and nodule antioxidant activities. Physiol Plant 2005; 124: 4–11.

86. Entcheva P, Phillips DA, Streit WR. Functional analysis of Sinorhizobium meliloti genes involved in biotin synthesis and transport. Appl Environ Microbiol 2002; 68: 2843–2848.

87. López-Maury L, Marguerat S, Bähler J. Tuning gene expression to changing environments: from rapid responses to evolutionary adaptation. Nat Rev Genet 2008; 9: 583–93.

88. Whitehead A, Crawford DL. Variation within and among species in gene expressionL: raw material for evolution. Mol Ecol 2006; 15: 1197–1211.

89. diCenzo GC, Benedict AB, Fondi M, Walker GC, Finan TM, Mengoni A, et al. Robustness encoded across essential and accessory replicons of the ecologically versatile bacterium Sinorhizobium meliloti. PLoS Genet 2018; 14.

90. diCenzo GC, Wellappili D, Golding GB, Finan TM. Inter-replicon Gene Flow Contributes to Transcriptional Integration in the *Sinorhizobium meliloti* Multipartite Genome. G3: Genes|Genomes|Genetics 2018; 8: g3.300405.2017.

91. Bell GDM, Kane NC, Rieseberg LH, Adams KL. RNA-Seq Analysis of Allele-Specific Expression, Hybrid Effects, and Regulatory Divergence in Hybrids Compared with Their Parents from Natural Populations. Genome Biol Evol 2013; 5: 1309–1323.

92. Hochholdinger F, Hoecker N. Towards the molecular basis of heterosis. Trends Plant Sci 2007; 12: 427–432.

