## Supplemental Material file S3 for "Dissecting transcriptomic signatures of genotype x genotype interactions during the initiation of plant-rhizobium symbiosis": Supplemental Table S1 (List of strains and alfalfa cultivars).docx

**Supplemental Table S1. List of strains A) and alfalfa cultivars B)**

**A)**

| **Strain** | **Host plant of original isolation** | **Genome sequence** | **Reference** |
| --- | --- | --- | --- |
| Rm1021 | *M. sativa* | GCA_000006965.1 | [1] |
| AK83* (DSM23914) | *M. falcata* | GCA_000147795.3 | [2] |
| BL225C (DSM23913) | *M. sativa* | GCA_000147775.3 | [2] |

* AK83 strain is also present, as original specimen after initial isolation, in the culture collection of All-Russia Institute of Agri- cultural Microbiology (RIAM, St. Petersburg, Russia).

**B)**

| **Name of the cultivar** | **Germplasm type** | **Fall dormancy** | **Producer of the material** |
| --- | --- | --- | --- |
| Camporegio | *Medicago x varia M. falcata* | 3 | CREA-FLC, Lodi, Italy |
| Verbena | *M. falcata* | 4 | CREA-FLC, Lodi, Italy |
| Lodi | *M. sativa* | 6 | CREA-FLC, Lodi, Italy |

1	Meade, H.M. *et al.* (1982) Physical and genetic characterization of symbiotic and auxotrophic mutants of Rhizobium meliloti induced by transposon Tn5 mutagenesis. *J. Bacteriol.* 149, 114 LP – 122

2	Galardini, M. *et al.* (2011) Exploring the symbiotic pangenome of the nitrogen-fixing bacterium Sinorhizobium meliloti. *BMC Genomics* 12, 235
