## Supplemental Material file S3 for "Dissecting transcriptomic signatures of genotype x genotype interactions during the initiation of plant-rhizobium symbiosis": Supplemental Table S2 (Primers used in this study).docx

| Gene name | Locus tag | Strain | Forward sequence (5’-3’) | Reverse sequence (5’-3’) |
| --- | --- | --- | --- | --- |
| *rplM* | SMc01804 | Rm1021 | AAGCGGCCTTCGATGATCTG | CTCCACCGGCAGAAGTACAC |
| *nodA* | Sma0869 | Rm1021 | ACCACCAGGAGCTCTCAGAA | TATCCCGACCGAGTCGTAAG |
| *nodB* | Sma0868 | Rm1021 | TGAGATTGTCGAGGCAAGTG | GGATCTGCCGACCAATGTAT |
| *sinI* | SMc00168 | Rm1021 | TCCCGAAATCTCCCAGGATC | ACCGTGACGATATGGCTGAT |
| *emrB* | SMc03167 | Rm1021 | AATGCGTCGGGACTCTACAA | GCTGCGTGAGGATAGTGTTG |
| *fixO2* | SMa0766 | Rm1021 | GTTTCAGTGGGGATCGAAGC | GAGGTTCGATCATGTGCTGC |
|  | SinmeB_6271 | BL225C | ACAATCTTTTCGTCGCAGCC | CGAACAAGATCCGCCACAAT |
|  | SinmeB_4750 | BL225C | TATGTCGTCGGCCAGATCC | GCCGGTATAGCTTCCCTTGA |
|  | Sinme_4706 | AK83 | CGAAGAGAAGACCAAAGGCG | CTGGATCTGACCCTTTTCGC |
|  | Sinme_6882 | AK83 | GTAATGTGACGACGCCCTTC | ACGTTCCATTTGTCGGCTTT |
