## Supplemental Material file S3 for "Dissecting transcriptomic signatures of genotype x genotype interactions during the initiation of plant-rhizobium symbiosis": Supplemental Table S4 (Metabolites mostly differing among root exudates).docx

**Table S4**. Metabolites with the greatest differences among root exudates. Results of Simper analysis based on the decomposition of the Bray-Curtis dissimilarity obtained from each peak ID value. The peak ID, the Average dissimilarity, the percentage of contribution, and hypothetical compound after a search of the PubChem library (<https://pubchem.ncbi.nlm.nih.gov>) are reported.

**Camporegio vs. Lodi**

| **Peak ID** | **Av. dissim** | **Contrib. %** | **Hypothetical compound** |
| --- | --- | --- | --- |
| PP_11707 | 21.08 | 29.16 | N-Acetyl-L-leucine |
| PP_16427 | 8.518 | 11.79 | [DL-Tryptophan](https://pubchem.ncbi.nlm.nih.gov/compound/1148) |
| PP_01693 | 5.773 | 7.988 | [Cytosine](https://pubchem.ncbi.nlm.nih.gov/compound/597) |

**Camporegio vs. Verbena**

| **Peak ID** | **Av. dissim** | **Contrib. %** | **Hypothetical compound** |
| --- | --- | --- | --- |
| PP_11707 | 17.08 | 24.3 | [N-Acetyl-L-leucine](http://www.chemspider.com/Chemical-Structure.64075.html) |
| PP_16427 | 6.941 | 9.876 | [DL-Tryptophan](https://pubchem.ncbi.nlm.nih.gov/compound/1148) |
| PP_01693 | 4.187 | 5.957 | [Cytosine](https://pubchem.ncbi.nlm.nih.gov/compound/597) |

**Lodi vs. Verbena**

| **Peak ID** | **Av. dissim** | **Contrib. %** | **Hypothetical compound** |
| --- | --- | --- | --- |
| PP_13281 | 10.57 | 21.59 | [3,5-Dihydroxyphenylglycine](https://pubchem.ncbi.nlm.nih.gov/compound/108001) |
| PP_14042 | 2.207 | 4.508 | [Val-Ala](https://pubchem.ncbi.nlm.nih.gov/compound/334517) |
| PP_11707 | 1.995 | 4.075 | [N-Acetyl-L-leucine](http://www.chemspider.com/Chemical-Structure.64075.html) |
