## Supplemental Material file S3 for "Dissecting transcriptomic signatures of genotype x genotype interactions during the initiation of plant-rhizobium symbiosis": Supplemental Table S5 (CHN).docx

**Table S5. Chemical composition of root exudates**. The percentage of nitrogen (N), carbon (C) and hydrogen (H) are reported for the root exudates of the three cultivars and the blank (medium stored in the apparatus used to collect root exudates). Sulphur was not detected in any sample and is omitted from the table.

|  | **N%** | **C%** | **H%** |
| --- | --- | --- | --- |
| **Blank** | 0 | 0.0172 | 0.49 |
| **Lodi** | 0.017 | 0.2458 | 56.61 |
| **Verbena** | 0 | 0.0601 | 55.47 |
| **Camporegio** | 0 | 0.0556 | 30.05 |
