## Supplementary figures and images for "Dissecting transcriptomic signatures of genotype x genotype interactions during the initiation of plant-rhizobium symbiosis"

### Figure S1.png

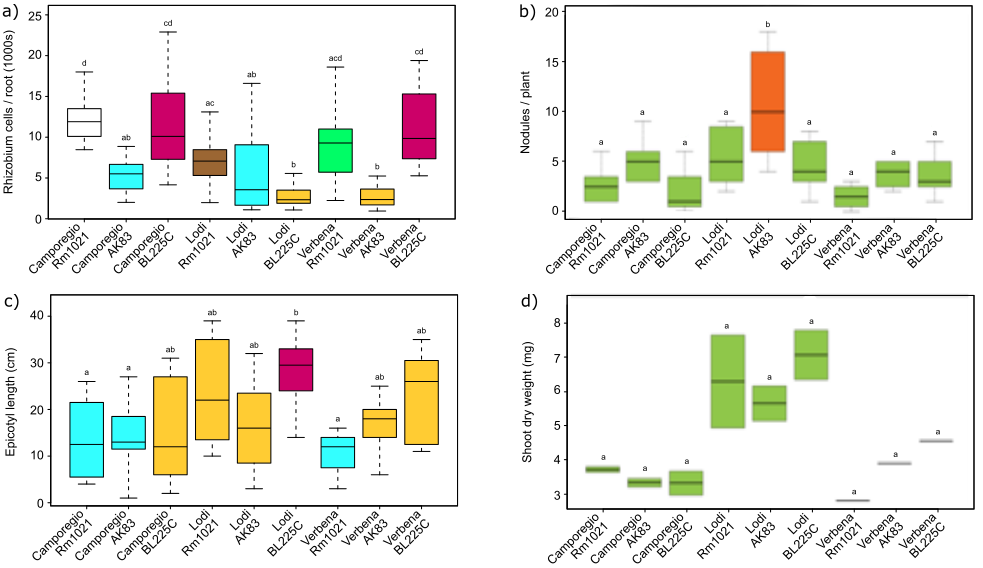

### Figure S2.png

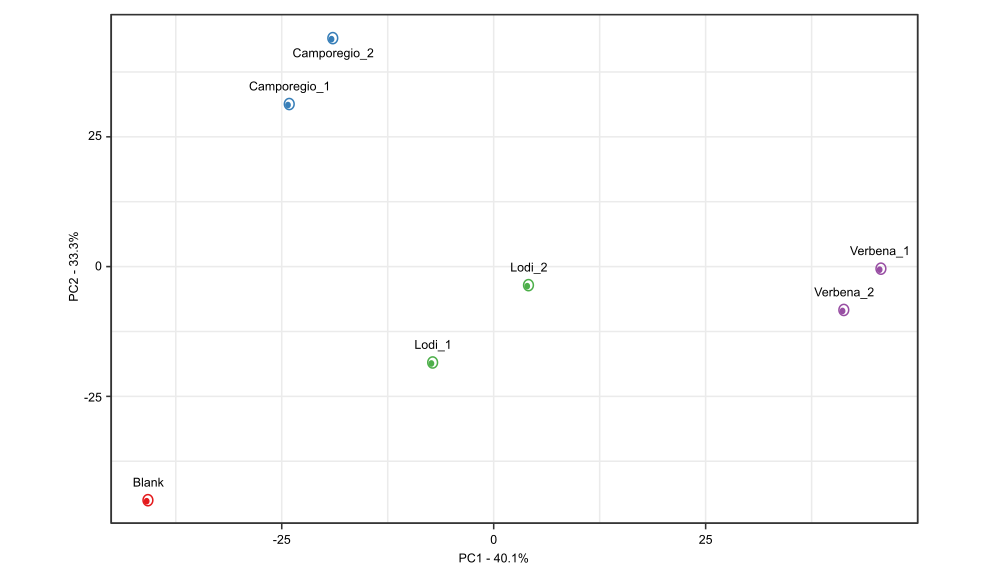
